# Integration of large, complex single-cell datasets with Harmony2

**DOI:** 10.64898/2026.03.16.711825

**Authors:** Nikolaos Patikas, Hongcheng Yao, Roopa Madhu, Soumya Raychaudhuri, Martin Hemberg, Ilya Korsunsky

## Abstract

Integrating single cell RNA-seq profiles is posing new challenges as datasets are rapidly expanding, now with over 100 million cells in the public domain. We present the latest version of the Harmony integration software, which efficiently scales to >100M cells and >1K datasets without specialized hardware. Moreover, optimizations to the underlying algorithm help prevent overintegration in biologically heterogeneous datasets. Harmony2 enables efficient, accurate integration of large, complex single-cell atlases.

## Introduction

Single-cell RNA sequencing has become a foundational assay for profiling cell types and states across healthy and diseased human tissues, with over 100 million profiles from more than 10,000 donors now publicly available^1–3^. Integrating these datasets into coherent reference maps requires computational algorithms that remove technical variation while preserving biological structure. As atlases grow in scale and heterogeneity, integration must balance two competing objectives: aligning shared cell states across batches while maintaining genuine biological differences.

Integration methods commonly face opposing failure modes. Underintegration leaves technical variation uncorrected, whereas overintegration merges biologically distinct cell types or states, particularly in heterogeneous datasets with non-overlapping populations. A range of approaches have been proposed, including Seurat^4^ CCA- and rPCA-based integration, matrix factorization frameworks such as LIGER^5^, regression-based correction methods such as ComBat-seq^6^, deep generative models such as scVI^7^, and regression-based embedding approaches such as Harmony^8^. While these methods have advanced the field, balancing batch correction with preservation of biological structure remains difficult. These challenges become particularly acute at the atlas scale where diverse data sets may have variable quality and may represent different cell types. At the same time, large datasets introduce substantial computational demands, requiring methods to scale efficiently with both the number of cells and the number of batches.

Harmony^8^ has been widely adopted for its speed and accuracy^9–11^. However, scaling to modern atlas-scale datasets requires improvements in both computational efficiency and robustness to heterogeneous batch structure. Here we introduce Harmony2, a redesigned implementation that addresses these challenges. Harmony2 incorporates optimized data structures, including a hybrid sparse–dense matrix backend and closed-form inversion for arrowhead-structured regression problems, enabling linear scaling in both cells and batches. To improve robustness in heterogeneous settings, Harmony2 introduces automated batch pruning and dynamic parameter tuning that reduce the influence of outliers or non-overlapping populations. Across large-scale benchmarks and real-world atlases, we show that Harmony2 scales efficiently, mitigates both over-and underintegration, and enables improved detection of rare and disease-associated cell populations.

## Results

### Harmony2 scales efficiently with cells and batches

We optimized the Harmony algorithm to scale efficiently with increasing numbers of cells and batches, while maintaining the core functionality (see **Methods**). Key optimizations include a sparse matrix backend that avoids redundant computation as batches increase, a closed-form solution to the regression step that replaces costly matrix inversion, and automatic pruning of batches with insufficient representation in each cluster. To evaluate our new implementation, we performed a series of benchmarks on the Tahoe-100M dataset^3^, comprising 1,135 batches (each batch corresponds to a well in one of 14 96-well plates) from 47 human cell lines cultured as mosaic cell line villages, totalling ∼100 million cells (**Fig S1a**).

Initially, we profiled how runtime and memory usage scale with increasing numbers of cells and batches, compared with the original Harmony1^8^. We downsampled the Tahoe-100M dataset to construct two benchmarking conditions using default Harmony parameters: (i) a fixed number of 1 million total cells with an increasing number of batches and (ii) a fixed number of 800 batches with an increasing number of cells. Harmony2 integrated 1M cells from 800 batches in less than 1 minute on the CPU, achieving a 203-fold speedup over the old version and a 12.5-fold reduction in memory usage. Importantly, our benchmarks show that the speedup increases proportionally to the number of batches. Using 1M cells and increasing the number of batches from 50 to 800 has no impact on runtime or memory usage for Harmony2 (**Fig 1a,1b**). In contrast, Harmony1 showed a linear overhead by the inclusion of additional batches (**Fig 1a,1b**). When fixing the number of batches and increasing the number of cells from 1 to 16 million, both versions of Harmony showed linear scaling of runtime and memory (**Fig 1c,d**), with Harmony2 showing much more favorable coefficients. Harmony2 required only 2.1 GB and 1.3 minutes per million cells, while Harmony1 required an additional 37 GB and 43 minutes per million cells, failing to integrate more than 4 million cells (**Fig 1c,d**). Taken together, a simultaneous increase in batches and cells would cause Harmony1 to scale quadratically, whereas Harmony2 scales linearly.

**Figure 1.**
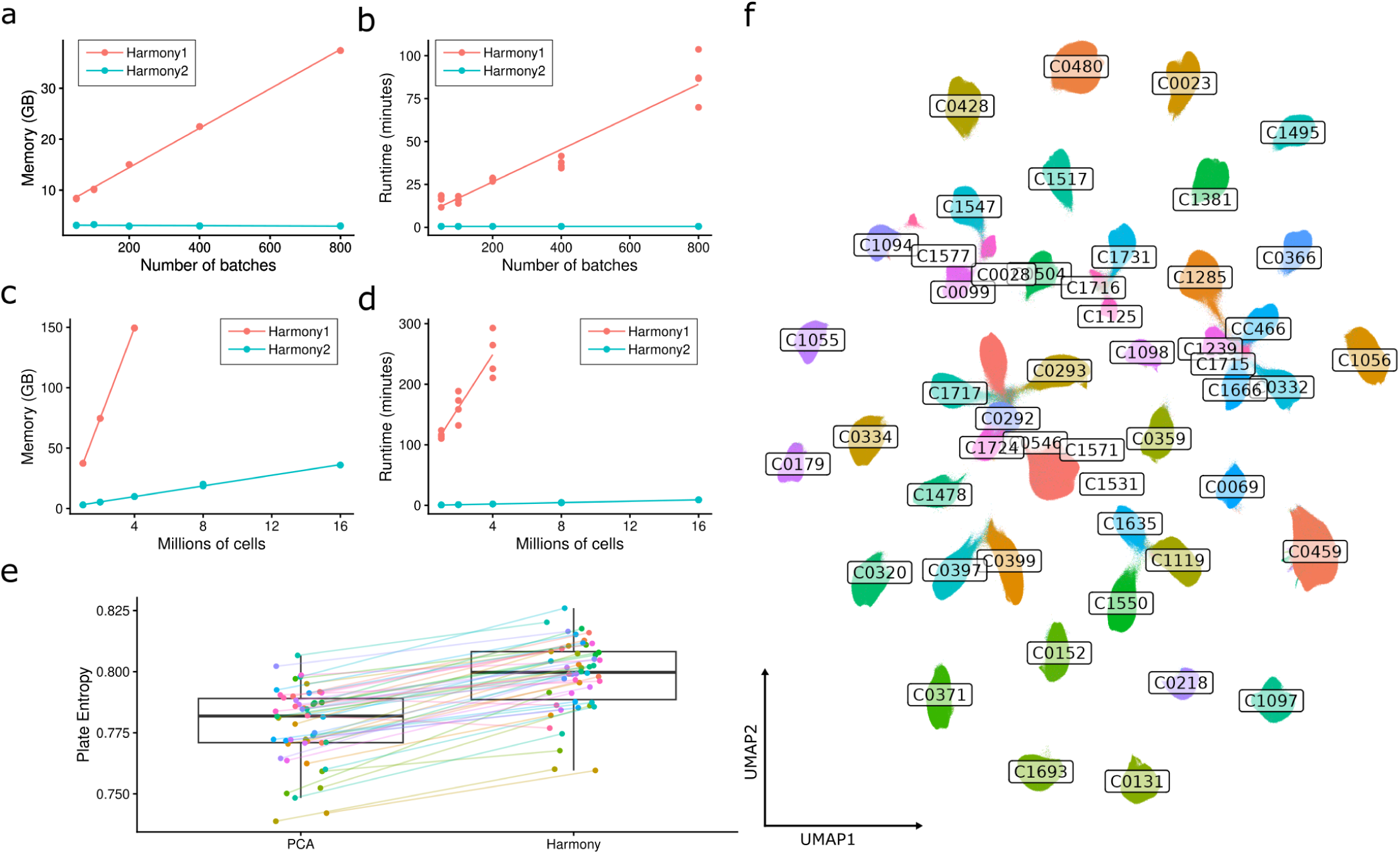
Harmony2 scales efficiently with cells and batches while preserving integration quality. (a,b) Memory (a) and runtime (b) performance of Harmony1 and Harmony2 using 1 million cells with an increasing number of batches. Lines represent linear regression fits to the observed data points. (c,d) Memory (c) and runtime (d) performance using 800 batches with an increasing number of cells (in millions). Lines represent linear regression fits. (e) Boxplot showing normalized single-cell plate entropy in the full Tahoe-100M dataset before (PCA) and after Harmony2 integration. Points represent per–cell line averages, illustrating the increase in plate mixing after integration. (f) UMAP embedding of the Tahoe-100M dataset after Harmony2 integration, colored by the 47 cell lines. A subset of 10 million cells sampled from the full Harmony-corrected embedding was used for UMAP computation and visualization. Labels indicate cell line identifiers positioned at the centroid of each cell line cluster.

We also assessed whether these performance gains affected integration quality. Both versions preserved biological separation between cell lines, with silhouette scores comparable to those of uncorrected PCA (**Fig S1b,c**). However, Harmony1 batch correction became less effective as the number of cells per batch increased, showing minimal improvement over PCA at 4M cells, and at 100 and 200 batches, whereas Harmony2 consistently improved batch entropy above PCA across all conditions (**Fig S1d,e**). Thus, Harmony2 delivers both substantial computational gains and more consistent batch correction across dataset configurations.

Finally, we sought to integrate the full Tahoe-100M dataset. Here, a successful integration would mix cells from different batches and plates, keeping cells from distinct cell lines separate. We quantify batch and plate mixing using normalized entropy on the kNN graph of Harmonized embeddings, and we quantify cell line separation using silhouette scores (see **Methods**). Harmony2 managed to integrate the dataset in 5.5 hours with approximately 233GB peak memory usage and 16 CPU threads. To assess the integration quality, we generated integrated UMAP embeddings from a subset of 10 million cells, which clustered primarily by cell line both before (**Fig S1f**) and after Harmony2 (**Fig 1f**). Using the batch-corrected Harmony2 embeddings computed on all cells, we observed a marked increase in single-cell plate entropy across all cell lines (**Fig 1e**) and across plates (**Fig S1g**), indicating that the original dataset exhibited moderate plate-driven batch effects that were effectively mitigated by Harmony2. We also observed a small increase in cell line silhouette score after Harmony2 integration, indicating slightly improved separation of cells by cell line in the latent space (**Fig S1h**). Overall, our findings show that Harmony2 demonstrates tractable scalability as the number of cells, batches, and conditions increases in complex datasets.

### Harmony2 balances batch integration and cell lineage preservation

To directly evaluate overintegration, where integration methods erroneously merge biologically distinct cell types (**Fig 2a**), we designed a controlled experiment with a predefined ground truth. We started from a single-cell atlas of inflamed human joint tissue comprising approximately 327,710 cells across 82 patient-specific batches and split the data into two imbalanced, non-overlapping groups of 41 samples each (**Fig 2b**). Group (i) contained only T cells, NK cells, and endothelial cells (n = ∼68,000), and Group (ii) contained only B and plasma cells, myeloid cells, and fibroblasts (n = ∼85,000) (see **Methods**). The two groups shared no cell types. This absence of overlapping lineages creates a stringent stress test, as methods can easily and incorrectly align related populations across groups, such as merging T and B cells. In this setting, a correct integration method should increase cross-batch mixing while preserving separation between distinct lineages. We quantified these using entropy-based metrics for batch mixing and cell type purity (see **Methods**), which both range from 0 to 1, with higher values for better mixing and stronger lineage preservation. We defined lineage labels by unsupervised clustering on the full, uncorrected dataset before splitting the data and without applying integration, ensuring that reductions in purity directly reflect overintegration rather than annotation uncertainty (**Fig S2a,b, Methods**). We benchmarked Harmony2 against Harmony1, PCA (no integration), and five widely used methods: scVI^7^, ComBat-seq^6^, Seurat-RPCA^4^, LIGER-Quantile Normalization (LIGER-QN)^5^, and LIGER-Centroid Alignment (LIGER-CA)^5^. We did not include Seurat CCA because it did not scale to the dataset sizes used in this benchmark (see **Methods**).

**Figure 2.**
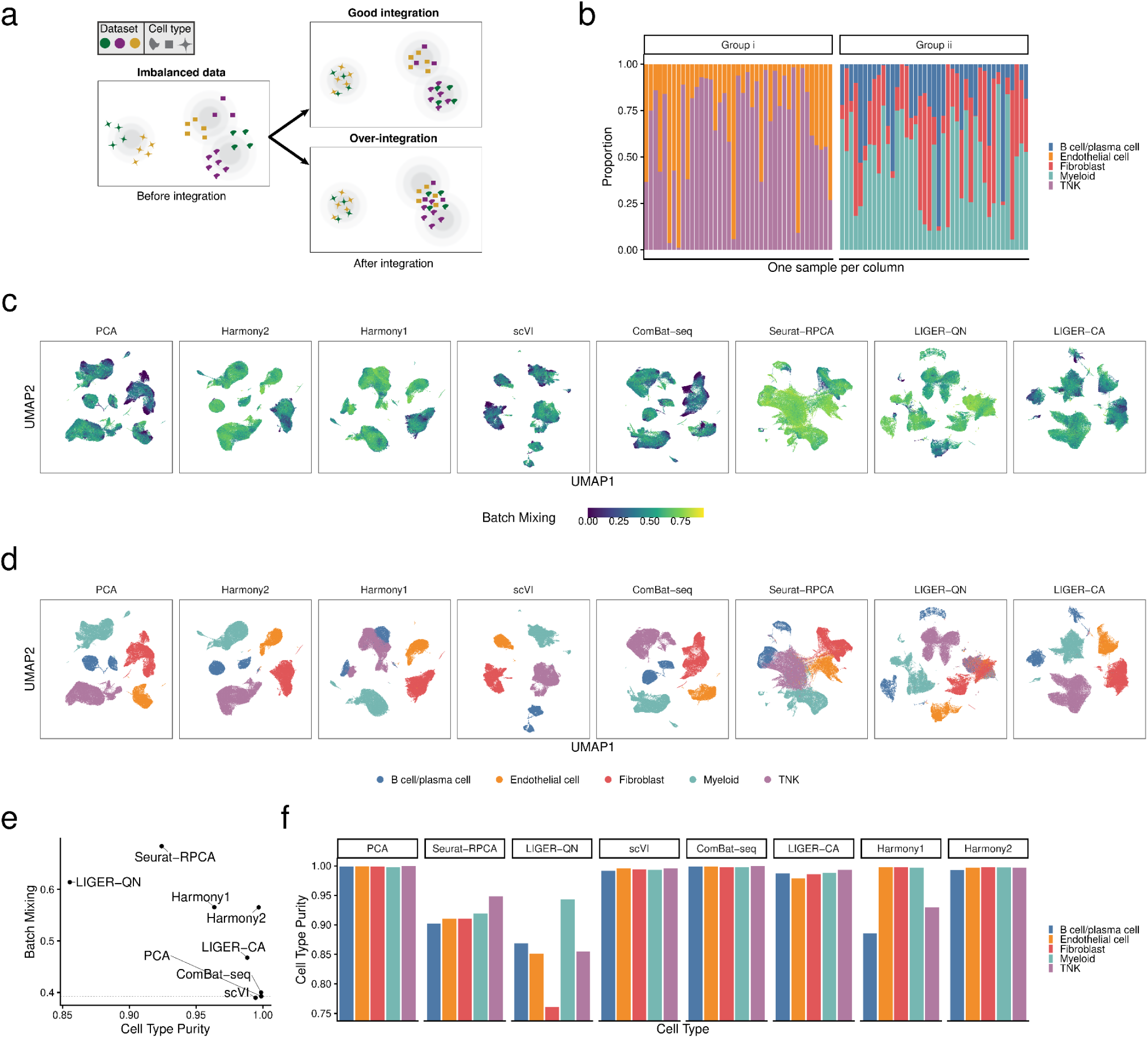
Controlled evaluation of batch mixing and lineage preservation in a heterogeneous integration setting. (a) Conceptual schematic illustrating integration outcomes in an imbalanced dataset with non-overlapping cell types across batches, highlighting correct integration versus overintegration. (b) Construction of the stress-test dataset from an inflamed human joint atlas, in which samples were divided into two imbalanced groups with no shared cell types; each column represents one sample and bars indicate the proportion of major cell types per sample. (c) UMAP embeddings after PCA or integration with the indicated methods, colored by batch mixing (local entropy). (d) The same UMAP embeddings colored by cell type. (e) Global comparison of batch mixing and cell type purity across methods, where batch mixing reflects cross-sample mixing and cell type purity reflects preservation of predefined lineage structure. (f) Lineage-specific cell type purity for each integration method across the five major cell types.

Comparisons of average batch mixing and cell type purity across methods revealed distinct integration regimes (**Fig 2c–e, Fig S2c**). UMAP visualizations reflected these differences, with PCA, scVI, and ComBat-seq showing limited batch correction, Seurat-RPCA and LIGER-QN exhibiting visible undesirable lineage mixing, and Harmony2 achieving strong batch correction while maintaining clear lineage boundaries (**Fig 2c,d, Fig S2c**). Quantitatively (**Fig 2e**), PCA set baseline expectations for batch mixing at 0.348, with minimal sample integration, but with high cell type purity at 0.999. scVI and ComBat-seq demonstrated performance comparable to this baseline ((0.346, 0.995) and (0.355, 0.999)), indicating minimal integration. LIGER-CA showed modest batch mixing with high purity (0.415, 0.989), reflecting limited mixing. In contrast, Seurat-RPCA and LIGER-QN substantially increased batch mixing but at the cost of reduced purity ((0.607, 0.928) and (0.545, 0.855)), consistent with aggressive overintegration. Harmony1 achieved intermediate batch mixing (0.502) with reduced purity (0.964). Harmony2 matched this batch mixing (0.502) while maintaining purity at 0.997, comparable to PCA.

Lineage-level analysis clarified these global patterns (**Fig 2f, Fig S2d**). Seurat-RPCA and LIGER-QN reduced purity across all lineages (0.903-0.948 for Seurat-RPCA and 0.762-0.943 for LIGER-QN), indicating widespread overintegration. In contrast, scVI and ComBat-seq maintained high purity ((0.992-0.996), and (0.998-1.00)) but often showed batch mixing at or below PCA levels, demonstrating underintegration. LIGER-CA exhibited modest batch mixing gains with small but consistent purity reductions across lineages (0.980-0.994). Harmony1 reduced purity primarily in B/plasma cells (0.886) and T/NK cells (0.930), the most transcriptionally related immune compartments, while largely preserving endothelial, fibroblast, and myeloid purity (0.997–0.999). Harmony2 preserved purity across all lineages (0.993-0.999) while maintaining elevated batch mixing (0.381-0.560). Together, these results show that Harmony2 increases batch mixing over PCA without sacrificing purity, whereas more aggressive methods (Seurat-RPCA and LIGER-QN) further increase batch mixing by trading away lineage separation, and more conservative methods (LIGER-CA, scVI, ComBat-seq) preserve purity by failing to integrate the batches. Unsurprisingly, most of the methods that offered the greatest sample integration, achieving the highest batch mixing, did so at the cost of overcorrection, compromising purity.

### Harmony2 powers unsupervised rare cell type detection

We hypothesized that Harmony2 would be able to scale to an extremely large data set, and preserve rare cell types by avoiding overintegration. We used Harmony2 to re-analyze the Human Lung Cell Atlas (HLCA)^12^, a collection of 2.3 million high-quality cells across 484 donors from 48 datasets. By pooling cells and donors, we expected to increase the power to detect rare (<1%) but functionally important epithelial cells: ionocytes, tuft cells, and neuroendocrine cells. Previous analyses relied on specialized labeling approaches, either supervised projection of disease cells onto a labeled atlas of healthy cells^12^ or supervised enrichment using a pre-defined gene signature^13^. We hypothesized that integrating all healthy and diseased cells with Harmony2 would group these rare epithelial populations, thereby eliminating the need for specialized strategies for rare cell type detection.

We used a hierarchical approach whereby we first integrated all 2,157,367 cells after QC to identify four major cell lineages with unsupervised clustering (**Fig 3a**). We then repeated Harmony2 integration and clustering on all 677,231 epithelial cells for two iterations (**Fig 3b,c**) and used the marker genes to match subclusters to rare cell types of interest (**Fig 3d,e**). We found four such clusters: one with neuroendocrine cells, one with ionocytes, one with mature tuft cells, and one with POU2F3^+^ tuft-ionocyte progenitors (TIP) (**Fig 3c,e**). For all rare cell types, we achieved high sensitivity and precision using previously published^13^ gene signature-based labels on healthy cells as ground truth (**Fig 3f,g**). Notably, we were able to identify twice as many mature tuft cells, which are reported to be very rare (∼0.002%^13^), compared to HLCA (n=37 vs 18, sensitivity= 0.698 vs 0.340) with high precision (0.88), with an additional 35 previously unlabeled mature tuft cells from 15 disease samples (**Fig 3f,g, Methods**).

**Figure 3.**
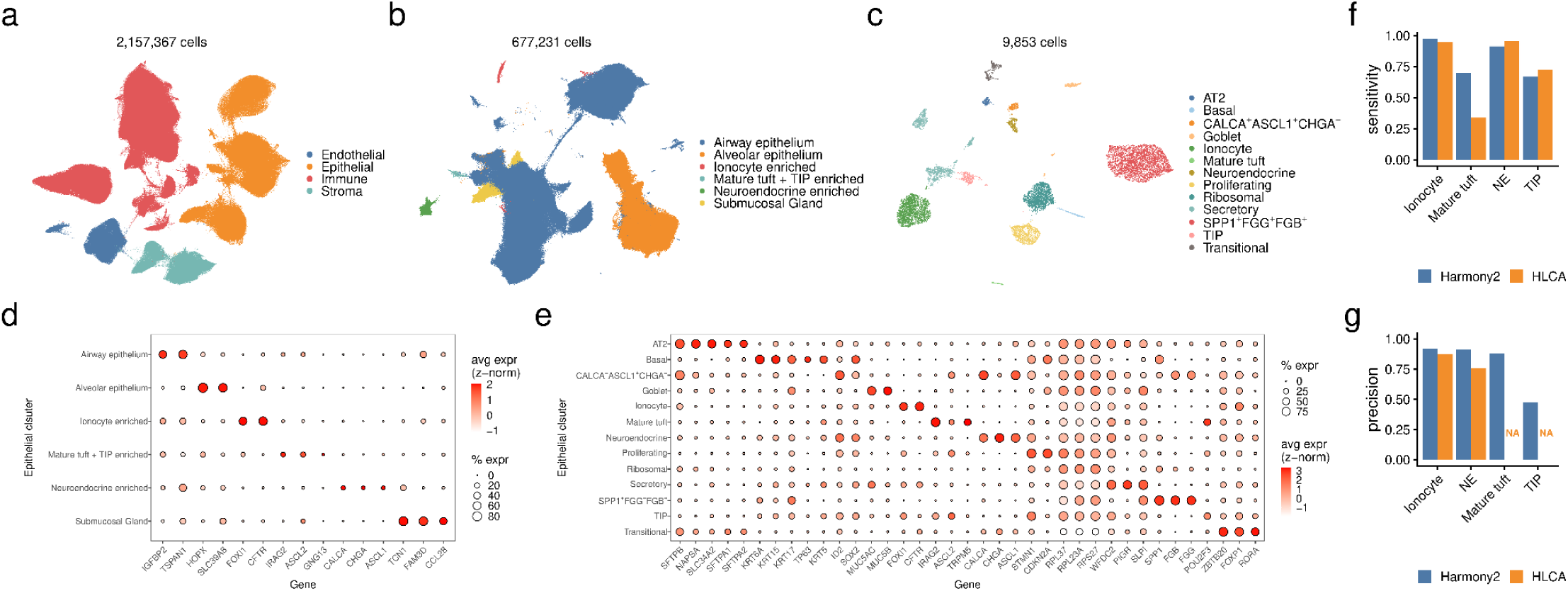
Harmony integration enables unsupervised identification of rare epithelial cell populations in the Human Lung Cell Atlas (HLCA). (a) UMAP embedding of all 2,157,367 quality-controlled HLCA cells after Harmony integration, colored by major cell lineage. (b) UMAP embedding of 677,231 epithelial cells following lineage-specific reintegration and clustering, highlighting major epithelial subtypes, including clusters enriched for ionocytes, neuroendocrine cells, and mature tuft cells with TIP enrichment. (c) UMAP embedding of 9,853 rare epithelial cells following focused reintegration, revealing ionocytes, neuroendocrine cells, mature tuft cells, TIPs, CALCA⁺ASCL1⁺CHGA⁻ cells, and additional epithelial subtypes. (d) Dot plot of marker gene expression across epithelial clusters. (e) Dot plot of marker gene expression across rare epithelial clusters; for (d) and (e), average log-normalized gene expression was computed per cluster and Z-normalized per gene, with dot size indicating the percentage of expressing cells and color indicating scaled average expression. (f,g) Sensitivity (f) and precision (g) of Harmony-derived labels and original HLCA labels for four rare epithelial cell types, using labels from Shah et al.^13^ as ground truth.

Additionally, we identified two clusters of previously unannotated epithelial cells derived from a small number of donors: CALCA⁺ASCL1⁺CHGA⁻ neuroendocrine-like cells and SPP1⁺FGG⁺FGB⁺ cells **(Fig. 3c,e**). We further focused on the CALCA⁺ASCL1⁺CHGA⁻ neuroendocrine-like cluster, as it closely resembles canonical neuroendocrine cells, and found that the majority of its cells were originally unannotated (79.9%) and predominantly derived from tumor samples (92%) from three patients^14,15^. By referring to the original study^14^, we concluded that this cluster consists of tumor cells expressing canonical neuroendocrine markers. Notably, CALCA⁺ASCL1⁺CHGA⁻ neuroendocrine-like cells were present at high frequency in one patient (69% of epithelial cells) who was diagnosed with both lung adenocarcinoma and large cell neuroendocrine carcinoma, but at low frequency in the other two patients (1% and 2% of epithelial cells), both of whom were diagnosed with lung adenocarcinoma only. These results demonstrate that Harmony2-based integration can identify a disease-associated cell type that occurs at very low frequency and in only a small number of patients, and can distinguish it from a related rare cell type by integrating data across all healthy and diseased samples.

## Discussion

Large single-cell atlases increasingly combine datasets across tissues, platforms, and disease contexts, resulting in substantial heterogeneity and partial or complete non-overlap of cell types across batches. In such settings, evaluating integration performance becomes inherently difficult because true lineage structure is unknown and many populations lack shared anchors across studies. These tasks are particularly sensitive to overintegration. Apparent improvements in batch mixing may therefore reflect either successful correction or unintended merging of biologically distinct populations, making over- and underintegration difficult to distinguish using post hoc inspection alone. To address this challenge, we designed a controlled stress test using a large human single-cell dataset in which lineage labels were defined on the full dataset prior to splitting and without applying integration. By constructing two imbalanced groups with non-overlapping cell types, we created a heterogeneous but realistic scenario in which biological preservation could be evaluated against a known reference without circularity. This design exposed a clear pattern across methods: approaches that maximized batch mixing frequently did so by collapsing distinct lineages, whereas methods that preserved lineage structure often failed to increase mixing beyond baseline levels. In contrast, Harmony2 increased batch mixing while maintaining lineage structure in a setting that mirrors the complexity of modern atlas-scale integration.

Our adaptation of Harmony for atlas-scale datasets presents enormous opportunities for researchers to combine and reuse the >100M scRNA-seq profiles existing in the public domain. If sufficient metadata is available, it may be feasible for some studies to rely on publicly available samples as healthy controls, which could reduce experimental costs by up to 50%. The ability to integrate at scale also opens up entirely new opportunities to assemble large, heterogeneous datasets. This will increase statistical power and enable studies with a scope beyond what the original experiments were designed for. For example, one could combine data from studies of different neurodegenerative diseases, such as Alzheimer’s disease^16^, amyotrophic lateral sclerosis^17^, Parkinson’s disease^18^, and frontal temporal dementia^19^. Acquiring these samples is both costly and time-consuming, and a comparative meta-study could help uncover common vulnerabilities and disease mechanisms.

The scale and efficiency of Harmony2 enable a dynamic view of large cell atlases, in which integration can be flexibly refocused on the subset of cells most relevant to a given biological question. Much as a city street map—rather than a world map—is required to navigate to a specific street address, resolving fine-grained cellular programs often requires re-integrating only the relevant populations at an appropriate resolution. For example, subtle differences between regulatory T cell subsets or exhaustion-associated transcriptional programs are unlikely to be well resolved when all cell types are jointly embedded, even if global integration appears successful. Existing reference-mapping approaches instead rely on static atlases whose resolution is fixed at construction, implicitly assuming that the appropriate reference already exists. Harmony2 reframes this workflow by making dynamic atlas re-integration practical at scale: users can efficiently re-integrate all relevant cells from a large atlas together with new samples, without reprocessing or reannotating existing data. This approach preserves contextual embedding while maximizing sensitivity to study-specific biological heterogeneity, offering a flexible alternative to static reference mapping as datasets continue to grow in size, scope, and diversity.

## Online Methods

### Summary of Harmony algorithm

Here is a glossary of terms used by Harmony:

- *d*: number of dimensions in PCA embedding
- *B*: number of batches
- *N*: total number of cells
- *N_b_*: total number of cells in batch *b*
- *C*: number of covariates (e.g. tissue, dataset, sample, technology)
- *K*: number of Harmony clusters
- *R*: Cluster assignment probability matrix (*N* x *K*)
- *O*: Observed number of cells for each cluster and batch (*K* x *B*)
- *E*: Expected number of cells for each cluster and batch (*K* x *B*)
- *Y*: Centroids (*d* x *K*)
- *Z*: Input embedding, to be corrected in Harmony (*d* x *N*)
- *Φ*: Design matrix (*B* x *N*)
- *Φ\**: Original design matrix with an intercept term: *Φ*\* = 1 || *Φ*.
- *θ*: Diversity penalty, hyperparameter
- _●_ *W*: Regression coefficients of dimension d, for batch b in cluster k. *W_kb_*∈ℝ^*d*^
- *λ*: Ridge regression sparsity penalty, hyperparameter.
- λ^: Empirically estimated values for *λ*, specific to each cluster and batch (*K* x *B*)
- α: Scaling factor for λ^

We provide a focused summary of the Harmony algorithm here to help the reader understand the improvements made. For a full description, we refer to the original paper^8^.

Harmony takes as input PCA embeddings in *d* dimensions for N cells, along with information about B batch assignments, encoded in a design matrix Φ, and returns embeddings with corrected batch relations. The algorithm iterates through two stages, clustering and linear regression, until convergence. The clustering uses a modified k-means with soft assignments, represented by a matrix *R,* which gives the probability of cell *i* being assigned to cluster *j*. The algorithm also includes terms in the objective function to maximize the diversity, defined as the number of different batches, contributing to each of the *K* clusters. Once the cluster centroids and assignments have been computed, we use ridge regression to update the corrected embeddings based on the cluster assignments, regressing out each of the *C* covariates.

### Sparse design matrices

The time and space complexity of linear algebra operations on sparse matrices depends on their sparsity. By definition, the batch-design matrix is a one-hot encoded matrix with exactly C+1 non-zero elements per column, where C is the number of batch covariates and an extra unit term for the intercept term of each cell. It is easy to see that adding more batch levels while keeping the number of covariates the same increases matrix sparsity. As such, making *Φ* sparse has no impact on the correctness of the algorithm, which learns the same correction factors and is much more efficient in terms of memory usage and runtime. Our implementation uses column-sparse compressed matrices for *Φ* and *Φ\**, enabling fast indexing along a single axis. During regression, we also need to use the transpose matrix *Φ^T^*. Since *Φ* and *Φ^T^*are constant through the Harmony algorithm, we also cache *Φ^T^*, avoiding repeated on-the-fly transposition of this matrix.

### Ridge-regression performance optimizations

#### Batch pruning

To calculate the batch correction factors in each harmony cluster *k*, the ridge regression step involves computing 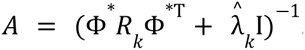, which requires the inversion of a (B+1)×(B+1) matrix. We leveraged the observation that each batch may contain cells in only a subset of Harmony clusters, and we only need to compute correction factors for those batches represented in the cluster. All other batches (by default, those with O_kb_/N_b_<10^-5^) are omitted, reducing the size of the matrix that needs to be inverted to solve the regression problem for that cluster. This batch-pruning step removes batches that have a very small fraction (by default, 1 in 100,000) of cells probabilistically assigned to the cluster, estimating correction factors only for batches with sufficient support.

The goal of this feature is two-fold. First, it alleviates the O(B^3^) cost of LU-based matrix inversions^20^ used by OpenBLAS^21^ by reducing the number of batches. When using a single covariate, this inversion is not necessary (see Arrowhead optimization), and the benefits are negligible; however, with more than one covariate, batch pruning yields substantial efficiency gains in the matrix inversion step. The second goal of batch pruning is to improve numerical stability. Low-support batches can make columns *A* linearly dependent for small λ^*_k_*, making the matrix non-full rank and non-invertible.

An important nuance emerges when we correct for multiple covariates. For instance, when we integrate over both dataset ID and sample ID (within a dataset), the dataset may contain cells from cell type *i,* but a specific sample may not. Here, Harmony2 prunes the sample from all clusters associated with cell type *i*, based on its probability assignment, and sets the sample-specific batch correction factors to 0 in these clusters. However, Harmony2 still learns dataset-specific batch correction factors for all clusters containing cell type *i,* and corrects the cells accordingly. The improvements from batch pruning are especially evident in large, heterogeneous datasets, where some cell types are missing from each dataset and therefore participate in only a subset of Harmony2 clusters.

#### Arrowhead optimization

We optimized the regression step for the common case in which a user specifies only one batch covariate (C=1) using arrowhead matrix inversion^22^. For an arrowhead matrix A with the following elements:

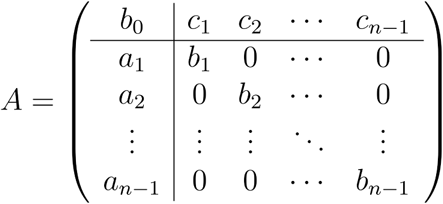

It has an efficient closed-form inversion formula:

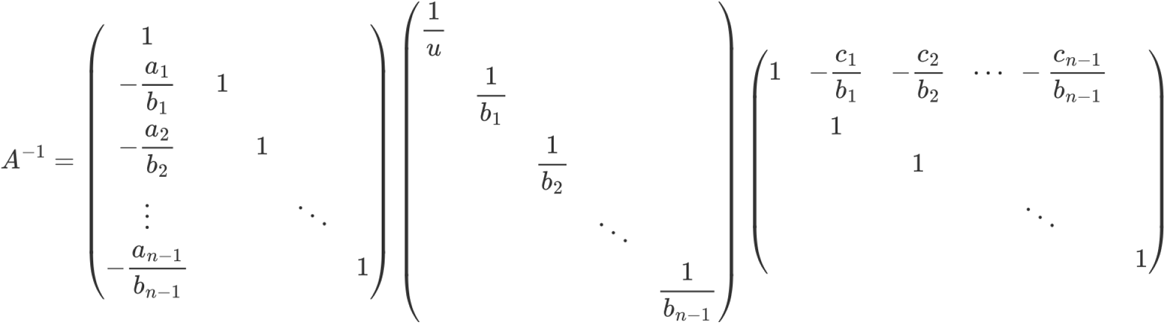

Rather than relying on the default LU factorization of a dense, unstructured matrix, which scales as O(B^3^), the closed-form arrowhead inversion exploits the matrix structure and can be solved in O(B) time. Harmony2 automatically detects when this optimization can be implemented and switches between default LU factorization and optimized arrowhead inversion as appropriate.

### Efficient k-means centroid initialization

Previously, the k-means centroids were seeded using the inefficient native R k-means implementation. We implemented a k-means++ based centroid seeding algorithm^23^ which requires O(KN) time and O(N) memory. Briefly, for each centroid *Y_k_*, we first select a random cell with embeddings *Z_i_* and compute the Euclidean distance between all other cells in the dataset and *Z_i_* to create a distance vector *D*∊ℝ^*N*−1^. We then choose the next centroid through weighted sampling of the remaining cells, choosing the next cell *j* with the lowest weight *s_j_ = log(u_j_)/D_j_*, with *u_i_* ∼ *U(0,1)*, the continuous uniform distribution from 0 to 1. If the selected centroid point has already been selected as the centroid for a different cluster, we select the cell *j* with the next lowest weight *s_j_*. This sampling makes it unlikely that we choose redundant cluster centroids that are close to one another.

### Stabilized diversity penalty

We improved the use of the cluster-diversity penalty *θ* to make the objective function more stable and decrease the risk of overcorrecting when *θ* is set incorrectly. The original algorithm had the following objective function.

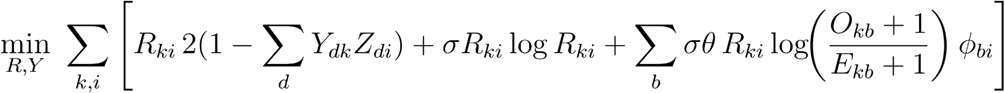

Here, the first term penalizes the cosine distance of points to their assigned centroids, the second maximizes the entropy of clusters (standard to soft k-means), and the third penalizes clusters with imbalanced batch composition (unique to Harmony clustering). In the last term, we compute the ratio of the number of cells from batch *b* in cluster *k* to the expected number under the independence assumption that every batch should have equal representation in every cluster. This ratio is minimized when *O_kb_*=*E_kb_* for every batch *b* and cluster *k*.

However, this ratio is numerically unstable when a batch lacks representation in a cluster (i.e. as *O_kb_* → 0). Our original solution of adding 1 to *O_kb_* and *E_kb_* fails when the total number of cells is large.

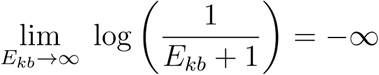

In practice, when *O_kb_* is small and the total number of cells is large, the magnitude of the diversity penalty becomes very large, dominates the objective function, and encourages diversity at the cost of high distances between cells and their centroids, which may lead to overintegrated clusters. We fixed this problem by reformulating the objective function to be scale invariant.

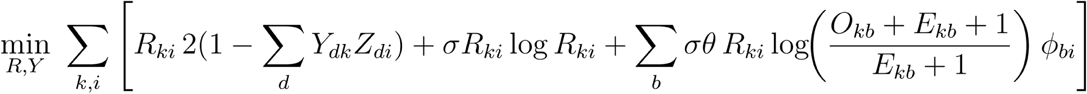

With this new function, the limit of the diversity penalty approaches log(1)=0, rather than − ∞.

We next re-derived the algorithm to minimize the new objective function. This change only impacts the soft cluster assignment computation - the E-step in the EM algorithm. Without showing the full derivation, here is the updated clustering assignment formula:

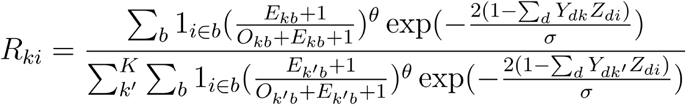

### Dynamic lambda estimation for controlling over-integration

During the ridge regression correction step, the penalty parameter *λ* controls the shrinkage of the regression coefficients, which is given by the vector W_kb_ for cluster *k*, batch *b*, which helps prevent over-integration.

Empirically, we observed that the shrinkage level for W_kb_ depends on the values of O_kb_ and λ. Specifically, if O_kb_<<λ, then ridge regression shrinks ||W_kb_|| to nearly 0. This situation occurs when cell *i* from batch *b* does not belong to cluster *k* but *R_ik_*>0, reflecting a soft assignment with negligible probability R_ik_. Therefore, λ must be large enough to be robust against these outlier soft-cluster assignments. At the same time, λ should be small enough to avoid shrinking regression coefficients of batches with small numbers of cells in a cluster.

In Harmony1, λ is fixed to 1, which is not always sufficient to shrink the regression coefficients of outlier batches to almost zero to avoid over-integration. Batch pruning, as mentioned above, can help by excluding some batches during the calculation of *W_k_*. However, the main purpose of batch pruning is to speed up calculations and to improve numerical stability. To prevent over-integration while accounting for data structure dynamically, we developed a dynamic approach for estimating the sparsity penalty with the following formula:

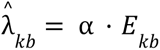

Based on empirical testing, a batch *b* for a cluster *k* typically requires more than (0.1 * *λ_kb_*) cell probability assignment to offset the shrinkage effect. Thus, we have found that α=0.2 can prevent undesired batches from being integrated into a cluster.

### Benchmarking

#### Runtime and memory profiling

The benchmarks were run using 20 principal components, which were subsets of the Tahoe-100M main dataset analysis. To compare the performance of the new and old Harmony, we used a shared high-performance infrastructure with the SLURM job manager, setting a 200GB per-run limit (350GB for the full Tahoe-100M dataset) using the exclusive flag. We ran four replicates per experimental condition, setting a different random seed for each. To profile Harmony’s runtime, we measured the RunHarmony() routine’s runtime using the peakRAM package. The most reliable way to measure memory usage was to probe the Linux process tree at /proc/<PROCESS-PID=/status for the VMPeak property, which records the peak memory a process requested in its lifecycle. To measure Harmony memory requirements, we measured VMPeak before invoking Harmony (baseline) and after running Harmony. The reported memory usage was the difference between the peak and baseline values. We ran our benchmarks so that the VMPeak property reflects the current memory usage before invoking Harmony (e.g., avoiding variable deallocation, matrix slicing, or invoking R’s garbage collector), ensuring the difference reflects Harmony’s memory footprint before the benchmark measurement. The coefficients reported were calculated by fitting linear models to the measurement data.

#### Single-cell entropy and silhouette scores

To assess the kNN graph homogeneity, we used the Normalized entropy calculation^10^, which yields values in [0,1] by normalizing using the maximum theoretical entropy. In the kNN graph, the maximum theoretical entropy is given by *log(min(K,#levels))*, where #levels is the number of distinct levels in a given covariate. For example, for k=20 neighbors, measuring the plate entropy where #plates is 14, the maximum theoretical entropy is log(14), and all Shannon entropies calculated are scaled by 1/log(14). In the Tahoe-100M dataset, the single-cell entropies were averaged over conditions. To assess the quality of the datasets, we used the simplified silhouette score^24^, which yields values in [-1, 1] for each cell. Then we summarized the differences per condition, and tracked the conditions before and after Harmony.

#### Tahoe-100M

##### Preprocessing

To analyse the Tahoe-100M dataset, we sourced the parquet files from HuggingFace at https://huggingface.co/datasets/tahoebio/Tahoe-100M and converted those to h5 and h5ad scanpy files^25^. Each sample consists of cells from a single condition (1 96-well plate) for a total of *N* samples. To process the data, we performed no filtering, resulting in 95,596,109 cells. The count data were normalized to 10k counts and log1p.

##### Feature selection

To select features in the Tahoe-100M dataset, we identified highly variable genes (HVGs) from *G* genes for each sample using scanpy’s default routine. In our samples, the HVGs varied substantially from 3,544 to 7,516. Subsequently, we performed a meta-analysis to determine a cutoff *L*,* and we consider a gene a feature if it is highly variable in at least *L\** of *n* samples. To determine *L\**, we defined the random variable X as a series of *N* Bernoulli trials with *p_i_=g_i_/n* where *g_i_* is the number of highly variable genes in sample *i*. If *p_i_=p* for *n* samples (all the samples have an identical number of HVGs), X ∼ *B(n,p),* where *B* is the binomial distribution. Using the normal approximation of the binomial distribution, we can see that X ∼ Ν(μ,σ^2^) has a *μ=np* and *σ^2^=np(1-p)*. Assuming *p_i_ = p*, we can factorize the equation for different p_i_ for each Bernoulli Trial: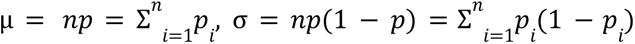 As such, we can expand the formulas and parameterize each p. Therefore, X can be approximated by the normal distribution 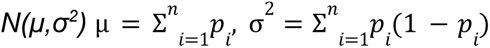 *L\** is given by *F*^−1^_*X*_ (*x*) = 0. 95 where *F^-1^* is the inverse normal cumulative distribution function. For the Tahoe-100M dataset, we have *n*=1,135 samples and this results in 6,578 HVGs.

##### Dimensionality Reduction

We calculated the eigenvectors by using an incremental principal components analysis approach from sklearn^26^. We achieve a substantial speedup by replacing the full SVD with a truncated SVD, setting the number of oversamples to 10. We streamed 1.5 million random cells from each sample in each SVD round to compute the final eigenvectors. After eigenvector calculation, we projected all cells to calculate the 50 most dominant Principal Components explaining a total variance of 24.3%.

##### Post PCA embedding processing

For the RunHarmony command, we selected the top 50 Principal Components with the default Harmony parameters, disabled batch pruning (each sample contains a similar mixture of cells), and set the number of threads to 16. For the kNN graph calculation, we used the approximate nearest neighbor method from the annoy^27^ Python package with 64 trees, and we computed the 20 nearest neighbors for the full dataset to generate the adjacency matrix. For the UMAP calculation, we downsampled the dataset to 10M cells, calculated the kNN using the same parameters to get the *G_20_* nearest neighbor graph adjacency matrix. Then, we cast *G_20_* as symmetric by applying the transformation *G_20_’ = G_20_ + G_20_ ^T^*, and we used the umap-learn^28^ package setting the min_dist=0.3.

#### AMP-RA

##### Cell type annotation

We collected all 327,710 cells, which were originally annotated with cell types from scRNA-seq data of AMP-RA. We removed proliferating cells, including “F-6: Proliferating sublining”, “T-18: Proliferating”, “NK-10: PCNA+ Proliferating”, “NK-11: MKI67+ Proliferating”, to simplify downstream analysis. The count matrix was log-normalized with a scaling factor of 10,000, and then the top 2000 HVGs were identified with function *vargenes_vst* from R package *singlecellmethods*. We then scaled the log-normalized matrix with only HVGs and performed PCA to calculate the first 30 PCs. We applied the PC matrix to *similarity_graph* function from R package *uwot* to generate the similarity graph, and then performed leiden clustering to annotate cell lineages, including T/NK cells, Fibroblast, Myeloid, Endothelial cell, B cell/plasma cell, which were referred to as cell type labels without batch correction.

##### Generation of the modified dataset

Based on cell type labels without batch correction, we randomly split samples into two groups and kept only corresponding cell types for each group: group (i) with 41 samples having only T, NK and endothelial cells, group (ii) with 41 samples having only B/plasma cells, myeloid cells and fibroblasts.

##### Harmony1 and 2

We log-normalized the count matrix with a scaling factor of 10,000, and then identified the top 2,000 HVGs. We then scaled the log-normalized matrix with only HVGs and performed PCA to calculate the first 30 PCs for downstream analysis. For both Harmony1 and Harmony2, we input the first 30 PCs and integrated over “sample” for 20 Harmony iterations, and turned off the early_stop argument, with all other parameters set to default. For Harmony2, the new theta penalty and dynamic lambda estimation were turned on.

##### Seurat-RPCA

R package *Seurat* version 4.4.0 was used to perform batch correction. Following the tutorial of *Seurat*, to correct for the batch variable of sample, we first split the data object to generate one data object per sample. We log-normalized the count matrix with a scaling factor of 10,000, and then identified the top 2,000 HVGs per sample individually. The global top 2,000 HVGs were selected by ranking genes according to the number of samples in which they were identified as HVGs. We then scaled the log-normalized matrix with only global HVGs and perform PCA to calculate the first 30 PCs per sample individually. We used *FindIntegrationAnchors* function from *Seurat* with reduction=”rpca” and dims=1:30, and the results were further supplied to *IntegrateData* function from *Seurat* with dims = 1:30, k.weight = 50, which generated a corrected log-normalized matrix. We next scaled the corrected log-normalized matrix and performed PCA to calculate the first 30 PCs as the final corrected embeddings.

##### scVI

Python package *scvi* version 1.4.1 was used to perform batch correction. Following the tutorial of *scvi*, to correct for the batch variable of sample, we first kept only genes with > 3 transcripts. We next performed highly variable gene selection with *highly_variable_genes* function from python package *scanpy* using flavor=“seurat_v3” and batch_key=“sample”. *scvi.model.SCVI.setup_anndata* and *scvi.model.SCVI* functions from *scvi* were used to set up model specifications with layer=“counts”, categorical_covariate_keys=[“sample”], n_latent=30. We acquired the final corrected embeddings by running *model.train* and *model.get_latent_representation* functions from *scvi*.

##### ComBat-seq

We log-normalized the count matrix with a scaling factor of 10,000, and then identified the top 2,000 HVGs. The count matrix with only HVGs was used as input for *ComBat_seq* function from R package *sva* version 3.35.2 with batch set as “sample”, which generated the corrected count matrix (for only HVGs). We then log-normalized the corrected count matrix with a scaling factor of 10,000. We next scaled the log-normalized matrix and performed PCA to calculate the first 30 PCs as the final corrected embeddings.

##### LIGER-quantile normalization and LIGER-centroid alignment

R package *rliger* version 2.2.1 was used to perform batch correction. Following the tutorial of *rliger*, to correct for the batch variable of sample, we first performed liger-specific normalization, HVGS and scaling with *normalize*, *selectGenes*, *scaleNotCenter* functions from *rliger*. We next performed integrative Non-negative Matrix Factorization using function *runIntegration* with k=30. We used the alignFactors function from *rliger* to perform factor alignment with method = “centroidAlign” for centroid alignment and method = “quantileNorm” for quantile normalization respectively, to generate the final corrected embeddings.

##### UMAP visualization

UMAP was generated using the *umap* function from *uwot* R package based on the embeddings. The parameters were set as: min_dist = 0.3, spread = 1.00, while all the other parameters were set as default.

##### Batch mixing and cell type purity

To characterize the level of batch mixing and cell type purity, we employed normalized entropy-based metrics. Normalized entropy for each cell is calculated as follows:

1. We construct the kNN graph for all cells based on corrected embeddings. k=50 was used for AMP-RA analysis.
2. For each cell, given a categorical variable A, normalized entropy was calculated as entropy / log(min(K, number of distinct levels of A))

For batch mixing, since the batch variable is “sample”, we calculated the normalized sample entropy for each cell and referred to it as the batch mixing. For cell type purity, we calculated 1 - normalized cell type entropy for each cell and referred to it as cell type purity. For the final batch mixing and cell type purity values per method or per method per cell type, average batch mixing and cell type purity across cells were calculated.

#### HLCA

##### Preprocessing and QC

To analyze scRNA-seq data of HLCA, we downloaded raw and normalized counts, cell type annotations, clinical and technical metadata of HLCA core and extension datasets from cellxgene (https://cellxgene.cziscience.com/collections/6f6d381a-7701-4781-935c-db10d30de293), which contains 2,282,447 cells from 48 datasets. We first performed QC to keep cells with number of unique genes >= 200, number of transcripts >=500 and <=50,000 and proportion of mitochondrial transcripts <= 0.1, which resulted in 2,157,367 cells.

##### Hierarchical Harmony integration and cell typing

To identify rare epithelial cells, we performed three iterations/levels of data normalization, PCA, Harmony integration, clustering, and cell typing. At all cells level, we log-normalized the count matrix with a scale factor of 10,000. Given the complexity and heterogeneity of data across different studies and datasets at this level, to identify highly variable genes, we first calculated standardized variance for each dataset independently using the “vargenes_vst” from R package “singlecellmethods”, and then computed the average standardized variance across datasets for each gene. We identified the top 2,000 genes with the highest average standardized variance as highly variable genes and then scaled the log-normalized matrix with only highly variable genes and performed PCA to calculate the first 40 PCs. We then input the first 40 PCs to Harmony2, and integrated over “sample” with theta=0 and all other parameters set to default. We then computed a similarity graph with the function “similarity_graph” from R package “uwot” and performed Leiden clustering using functions “from_adjacency” and “cluster_leiden” from R package “igraph”, with a resolution of 0.1. We labeled the cell type lineage based on marker genes and identified the epithelial cells. We then repeated this process at all epithelial cells level, and performed log normalization in the same way.

Since rare epithelial cells were not equally represented across datasets, we performed HVG selection in a slightly different way. We first calculated the standardized variance for each dataset independently as described above, and identified the top 2,000 genes with the highest standardized variance per dataset. For each gene, we then counted the number of datasets in which it appeared among the top 2,000 genes, and the 2,000 genes found in the highest number of datasets were selected as our HVGs at this level. We performed scaling and PCA in the same way and computed the first 100 PCs, using a larger number of PCs to enable the identification of small clusters. We then input the first 100 PCs to Harmony2, and integrated over “sample” with theta=0 and all other parameters set to default. We computed the similarity graph and performed Leiden clustering with a resolution of 0.8. Based on canonical marker genes of rare epithelial cells, we identified a cluster enriched in mature tuft+TIP, a cluster enriched in ionocytes, and a cluster enriched in neuroendocrine cells.

We repeated this process with only the three clusters enriched for rare epithelial cells. We log-normalized the count matrix in the same way and calculated standardized variance across all cells using the “vargenes_vst” function. We selected the top 2,000 genes with the highest standardized variance as the HVGs and performed scaling and PCA in the same way. We then input the first 40 PCs to Harmony2 and integrated over “sample” with theta=1 and all other parameters set to default. We computed the similarity graph and performed Leiden clustering in the same way using a resolution of 0.4, which allowed us to identify clusters representing ionocytes, neuroendocrine cells, mature tuft cells, TIPs, and CALCA⁺ASCL1⁺CHGA⁻ neuroendocrine-like cells, along with other cell subtypes.

##### Differentially expressed gene analysis

We performed differentially expressed gene analysis using the *wilcoxauc* function from R package presto^29^ with input of log-normalized gene expression and corresponding cluster labels.

##### UMAP

We generated UMAPs using the umap function from R package uwot. The parameters were set as: min_dist = 0.3, spread = 1.00, fast_sgd = TRUE, all the other parameters were set as default.

##### Accuracy

To get a high-confidence ground truth for rare epithelial cell type labels, we downloaded gene signature-based cell type labels annotated by Shah et al.^13^ from the github repository (https://github.com/TsankovLab/LungRareCells/tree/main/data).

There are four major rare epithelial cell types, including ionocytes, neuroendocrine cells, mature tuft cells, and TIPs. However, HLCA labels did not distinguish between mature tuft cells and TIPs, using only a tuft cells category. For the calculation of sensitivity, we assigned any mature tuft cells or TIPs labeled as tuft cells by HLCA to the category determined by Shah et al. However, we cannot use the same or similar approach when calculating precision, and thus the precision values of mature tuft cells and TIPs for HLCA were denoted as NA.

## Data and Code Availability

All data used in analyses for this paper will be deposited into Zenodo prior to publication. All code to produce results and figures will be deposited into the github repository on korsunskylab/harmonymanuscript2025. The Harmony2 software is available as an R package on github.com/immunogenomics/harmony and on CRAN.

## Supplementary Figures

**Supplementary Figure 1.**
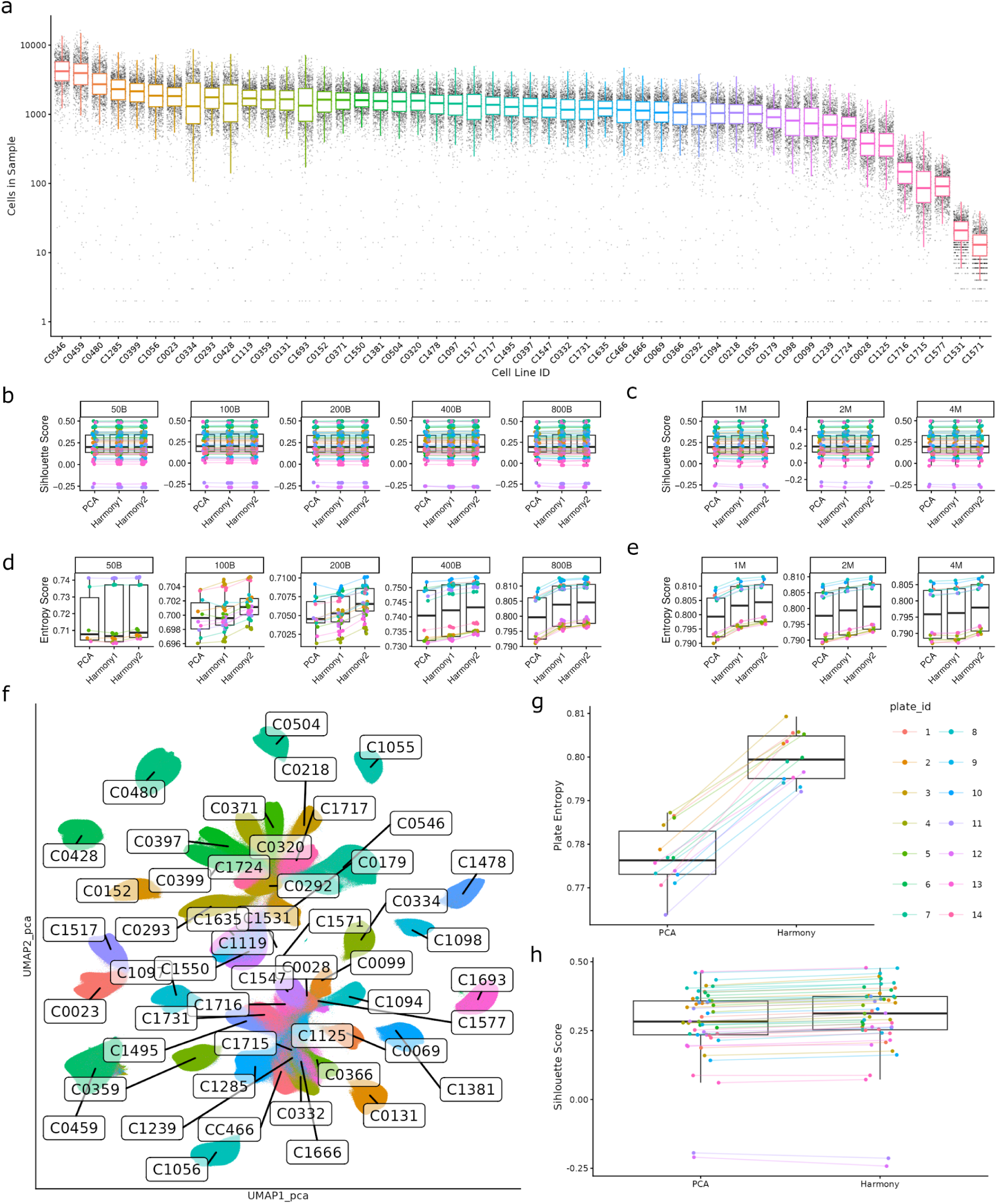
(a) Box plots showing the number of identified cells in the 1,135 samples in the Tahoe-100M dataset grouped by cell line. Each sample is expected to have a mixture of 47 cell lines and the cell line identity was demultiplexed using cell line genotype information. (b,c) Cell line silhouette scores for PCA, Harmony1, and Harmony2 across benchmarking conditions with increasing numbers of batches (b) and increasing numbers of cells (c). Each point represents the average silhouette score per cell line. (d,e) Normalized batch entropy scores for PCA, Harmony1, and Harmony2 across benchmarking conditions with increasing numbers of batches (d) and increasing numbers of cells (e). Each point represents the average entropy per cell line. (f) UMAP of the Tahoe-100M dataset using 50 principal components colored by the 47 cell lines. 10 million cells were sampled from the full principal component matrix and used for UMAP calculation and display. Labels indicate cell line identifiers positioned at each respective UMAP centroid. (g) Boxplot of normalized single-cell plate entropy in the full Tahoe-100M dataset before (PCA) and after Harmony2 integration. Points represent averages across plates, highlighting a consistent increase in plate entropy after integration. (h) Box plot of cell line silhouette scores before (PCA) and after Harmony2 integration of the full Tahoe-100M dataset. Each point represents the average single-cell silhouette score per cell line.

**Supplementary Figure 2.**
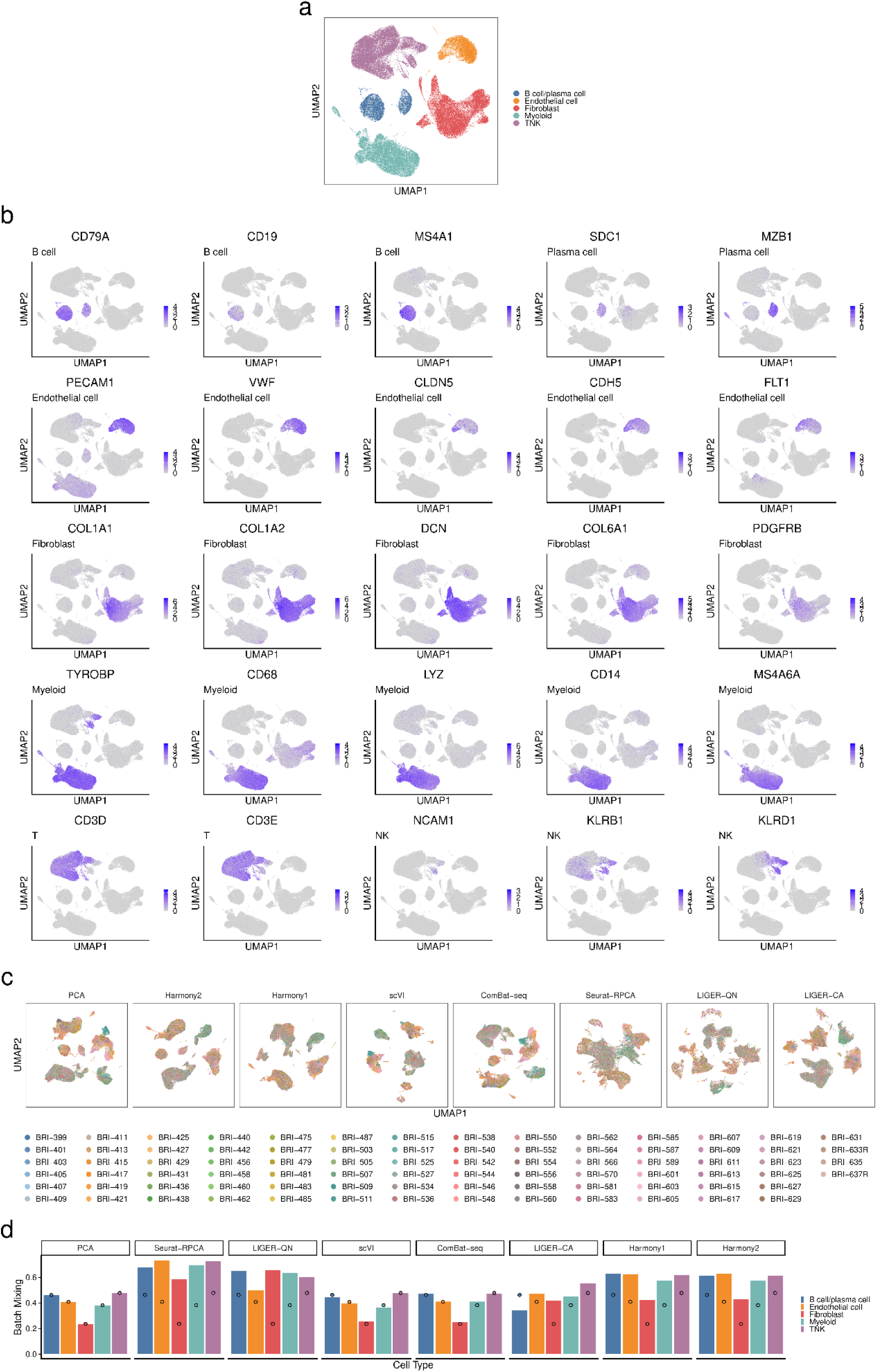
**(a)** UMAP for the unmodified full AMP dataset, colored by cell type. **(b)** UMAP for unmodified full AMP data, colored by marker gene log-normalized expression level. **(c)** UMAP embeddings after PCA or integration with the indicated methods for the modified AMP dataset, colored by sample. **(d)** Lineage-specific batch mixing for each integration method across the five major cell types. Dots correspond to the batch mixing levels of PCA, which are regarded as the baseline batch mixing levels.

## Acknowledgements

This work was supported by the National Institutes of Health (5K01AR078355, R03LM015261, R03OD039974, OT2OD033759) and the Chan Zuckerberg Foundation (Data Insights Grants #DI3-0000000372 and #DAF2022-249216). We thank members of the Korsunsky, Raychaudhuri, and Hemberg labs at BWH and the CZI Single Cell Science Program for insightful discussions.

## Author contributions

IK and MH conceived the research. NP and HY led software development of Harmony2 and jointly contributed to analyses. All authors participated in interpretation and writing the manuscript.

